# Operational lessons learned from simulating an elimination response to a transboundary animal disease in wild animals

**DOI:** 10.1101/2024.07.22.604666

**Authors:** K Chalkowski, KM Pepin, MJ Lavelle, RS Miller, J Fischer, VR Brown, M Glow, B Smith, S Cook, K Kohen, S Sherburne, H Smith, B Leland, KC VerCauteren, NP Snow

**Affiliations:** National Wildlife Research Center, Fort Collins, CO, 80521; National Wildlife Research Center, USDA-APHIS, Fort Collins, CO, 80521; Center for Epidemiology and Animal Health, United States Department of Agriculture, Fort Collins, Colorado, 80521; Center for Epidemiology and Animal Health, United States Department of Agriculture, Fort Collins, Colorado; National Feral Swine Damage Management Program, 4101 Laporte Avenue, Fort Collins, Colorado 80521, USA; National Wildlife Research Center, USDA-APHIS, Fort Collins, CO 80521, USA

**Author notes:** Corresponding author: Kayleigh Chalkowski. These authors contributed equally (K Chalkowski, KM Pepin).

## Abstract

Transboundary animal disease (TAD) introductions can have myriad economic, ecological, and societal impacts. When TADs are introduced into wild species, rapid and intense control efforts to reduce wild animal host populations are sometimes needed to eliminate the disease and prevent endemicity and spillover to domestic animal populations. Yet, such intensive efforts are non-trivial, and the rarity of TAD introductions means that personnel rarely have direct experience with these types of operations. Thus, explicit assessments of operational challenges for these kinds of efforts can provide direction to build emergency response preparedness capacity. Here, we simulated a TAD control effort in response to initial detection of a hypothetical index case of a TAD in wild pigs (*Sus scrofa*) (e.g., African swine fever; ASF). We used three removal methods (aerial control, trapping, and an experimental toxic bait). Then, we conducted an after-action assessment to identify operational challenges for rapidly reducing a population of invasive wild pigs within a simulated outbreak zone. We also simulated carcass recoveries of dispatched pigs, similar to what might be conducted during a response to a TAD with carcass-based transmission (e.g., ASF virus). Here, we describe operational challenges identified during our effort, alongside technological development solutions and *a priori* strategy needs to improve TAD response operation outcomes.

## Introduction

Transboundary animal disease (TAD) introductions to domestic animal populations can have far-reaching economic, ecological, and societal impacts (Chinchio et al., 2020; Paarlberg et al., 2008; Clemmons et al., 2021). When newly introduced diseases spillover from domestic animals into wild animal populations, ongoing transmission between wild and domestic animals can challenge disease elimination (Bengis et al., 2002, Pepin et al. 2023). As with any infectious disease (Steele et al. 2016), responses to TAD introductions in domestic wild animals both require rapid and effective detection and intense control efforts early in an outbreak to prevent spread and endemicity (Pepin et al., 2022, Bozzuto et al., 2020, Bolzoni et al., 2014). Despite the difficulty of these types of efforts, research addressing operational limitations of TAD emergency responses are limited.

TADs may become endemic in wildlife populations after an introduction (e.g., African swine fever in wild boar in eastern European countries (Chenais et al., 2019)), or they may spillover into wildlife species and rapidly fade out (e.g., foot-and-mouth disease virus in wild pigs (Pepin and Vercauteren, 2016)). Disease elimination can be achieved through a variety of strategies including wildlife culling or mass vaccination (Gortazar et al., 2015; Prentice et al., 2019; Nugent et al., 2015; Licoppe et al., 2023) to reduce the population of susceptible individuals below the minimum density threshold in order to prevent epidemic growth (Potapav et al., 2012). However, in cases where elimination is infeasible, other techniques including fencing or physical separation of wildlife from domestic animals, or ecological control strategies such as habitat manipulation, may be preferred methods for controlling TAD spillover to domestic animals or humans (Sokolow et al. 2019; Prentice et al., 2019; Gortazar et al. 2015).

Eliminating a newly introduced TAD in a wild animal population poses unique challenges compared to similar emergency responses, or wildlife management operations. TAD control in wildlife is different from responses to TADs in domestic animals because wild species are free-ranging, may be elusive, may reside in remote locations that are difficult to access, or have poorly characterized population densities, distributions, and movement behavior. Wildlife also may be subject to a variety of management and ownership policies that affect how control can be implemented in space and time. Finally, tools used for disease control in domestic animals (e.g., vaccination, population elimination) may not be efficacious or approved for use in wild species. Given these key differences, strategies for TAD control in domestic animals are not always transferable to TAD control in wildlife.

Elimination of TADs in wildlife is also operationally distinct from other kinds of wildlife management operations, such as damage mitigation. Compared to damage mitigation contexts, disease control operations require operational strategies grounded in disease ecology theory to minimize disease spread (Joseph et al. 2013). Specifically, this entails rapid, intensive deployment following an introduction (i.e., “hit early, hit hard”, Bozzuto et al., 2020; Pepin et al., 2022; Domenech et al., 2006) to prevent an outbreak; knowledge of behavioral responses to disease management to avoid counterproductive outcomes that facilitate disease spread (Prentice et al. 2019); intensive effort in a region over a long duration (i.e., longitudinal efforts (Gortazar et al. 2015)); and monitoring and interpretation of data with respect to disease dynamics (Gortazar et al. 2015).

Wild pigs (*Sus scrofa*), originating from a mixed ancestry of native wild boar and domestic feral pigs (Keiter et al. 2016), are widespread globally and can be infected by many transboundary animal disease pathogens including African swine fever virus (ASFv), classical swine fever virus, and foot-and-mouth disease virus (Miller et al., 2017; Meng et al., 2009; Ruiz-Fons et al., 2008), making them a high-risk wild species for potential establishment and persistence of TADs. Wild pigs are also one of the most widespread invasive species globally (Risch et al., 2021, VerCauteren et al. 2024), and inhabit much of North America (United States Department of Agriculture, 2023; Lewis et al., 2019). Accordingly, TAD response plans in the U.S. have incorporated response strategies for wild species, including wild pigs, alongside domestic livestock response plans (United States Department of Agriculture, 2020, 2023). These U.S. national strategic plans for responding to a TAD outbreak in wildlife, such as ASFv, involve large-scale culling actions that require rapid implementation, continuous intensive control, and iterative adjustment to current conditions (United States Department of Agriculture, 2020, 2023). Culling can be an appropriate and effective tool for controlling a TAD outbreak in invasive species (Gortazar et al., 2015), including wild pigs, but much of what is known about how to reduce wild pig populations comes from managing for population damage mitigation contexts rather than TAD eradication.

Previous efforts to eliminate newly introduced TADs such as ASFv in wild pigs and wild boar have found varying levels of success. Strategies that have been cited as drivers of successful ASFv elimination have included fencing to reduce movements, carcass recovery and removal, and intensive population reduction (Guberti et al. 2019, Palencia et al. 2023). Concurrently, many of these key strategies are frequently cited as high effort, costly, logistically or operationally challenging, and requiring highly trained personnel (Guberti et al. 2019). In fact, implementation challenges may in fact be a reason that some ASFv interventions utilizing these strategies have not been successful in eliminating the disease (Jo and Gortázar 2021). Thus, thorough consideration of operational challenges to implementing known disease elimination strategies, prior to an ASFv introduction, may improve national emergency response capacity.

Our objective was to identify operational challenges from an intensive reduction of a wild pig population within a simulated outbreak zone, and using common removal techniques for wild pigs (aerial operations, trapping, and an experimental toxic bait), alongside recoveries of carcasses dispatched by aerial operation and toxicant, similar to what may be conducted during a response to an ASF introduction (United States Department of Agriculture, 2023). We conducted an after-action assessment to identify important challenges and limitations of each control method that we tested in a TAD response context, as well as corresponding knowledge gaps, resource needs, and strategies to address the challenges we identified. Lastly, we discussed the findings of our after-action assessment in a TAD context beyond ASF and our study area.

## 2. Methods

### 2.1 Study site

We conducted this study in north-central Texas during Feb–May 2023. The study area (174.9 km^2^) was an active cattle rangeland, with variable wild pig density estimates ranging from 5-15 pigs/ km (Snow et al., 2024). The landscape was characterized as part of the south central semi-arid prairies ecoregion (level II), dominated by juniper (*Juniperus* spp.), mesquite (*Prosopis* spp.), scrub oak (*Quercus* spp.), and midgrass savanna (Bailey, 1980, 1998) bordered by croplands. During the study period, temperatures averaged 17.6 °C and a total 26.2 mm of precipitation occurred (National Oceanic and Atmospheric Administration, 2023). We chose three removal methods that are commonly used to reduce wild pig abundance: toxicant baiting, trapping, and aerial operations. We did conduct limited ground-shooting, but this method was inefficient in this habitat, and we did not have the personnel to thoroughly assess more than three removal methods. All research methods were approved by the USDA National Wildlife Research Center, Institutional Animal Care and Use Committee (QA-3311 and QA-3470).

### 2.2 Control procedures overview

Here, we describe a brief overview of our control methodologies to provide context for our observed operational challenges. More detailed descriptions of control methodologies are available in Snow et al. (2024), wherein we describe numbers of pigs removed with each technique alongside personnel effort, equipment and supply costs, and wild pig densities. For our removals, we divided the study area into 3 separate treatment zones, including aerial operations (46.3 km^2^), trapping (63.7 km^2^), and toxic baiting (64.9 km^2^)(Figure 1). Wild pig density estimates were variable across zones, with a higher density in the aerial zone compared to trapping and toxicant zones (∼10-15 pigs-km2, 5-12 pigs/km2, and ∼4-6 pigs/km2, respectively)(Snow et al 2024). Within the aerial and trapping zones, we designated 20.6 km^2^ and 18.4 km^2^ focal removal areas, respectively, where we conducted intensive control efforts to remove wild pigs (Figure 1). The location and size of these focal areas were designated based on known wild pig presence and our estimated ability to attempt removing wild pigs from the entirety of both areas. For toxic baiting, we were limited to 11 sites based on the amount of toxic bait we were approved to deploy in the US Environmental Protection Agency issued Experimental Use Permit (Reg. No. 56228-EUP-42).

**Figure 1.**
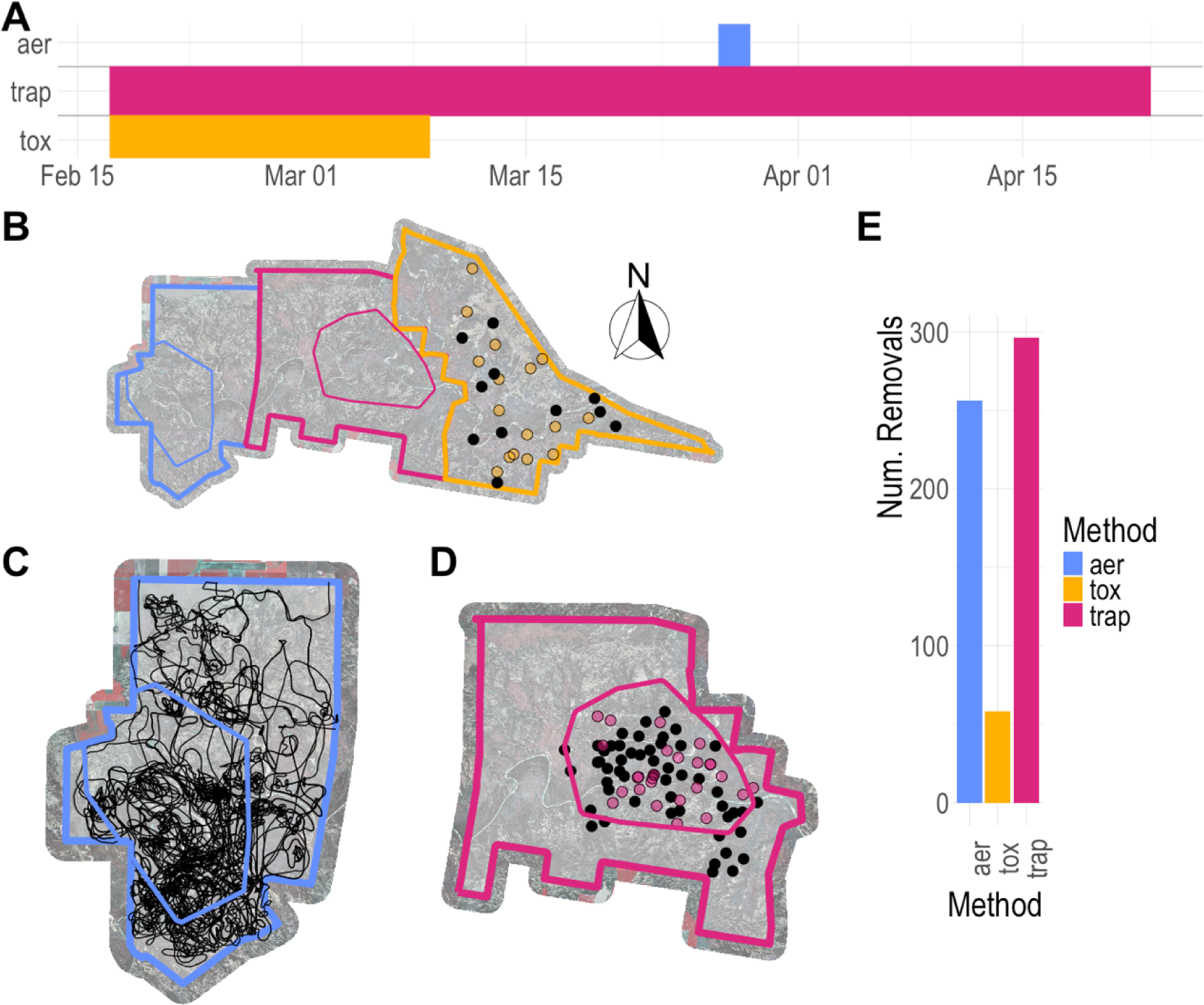
A. Timeline of activity for each wild pig (*Sus scrofa*) removal method, aerial operations were conducted from March 27-29^th^, 2023; toxicant operations were conducted from February 17-March 09, 2023; trapping operations were conducted from February 17-April 23, 2023. B. Study area in north-central Texas where wild pig removal activities were conducted as part of a simulated response to initial detection of a hypothetical index case of African swine fever virus. Outer polygons represent different study areas where removal activities were separately implemented. Inner polygons represent guidelines for where control activities should begin and spread outward, determined based on accessible terrain and/or known wild pig presence. Black dots indicate locations of 11 final toxicant deployment sites. Yellow dots indicate locations of sites bait sites to assess wild pig presence. Polygon zone colors: blue, aerial zone; magenta, trapping zone; yellow, toxicant zone. C. Aerial zone with inner polygons for initial guidelines, with black lines of flight paths for aerial control operation activities. D. Trap zone with inner polygons representing initial guidelines for where wild pig removals would take place, with black dots representing active trap sites (i.e., trap sites that were active for at least one trap night), and magenta dots representing bait sites to assess wild pig presence where traps were never deployed. E. Bar chart with numbers of wild pigs removed for each removal method.

### 2.3 Trapping Removals

We trapped at a total of 51 sites (Figure 1) during 76 days, totaling ∼728 trap nights. Generally, we used a whole-sounder approach for trapping, where we targeted entire groups of wild pigs (Lewis et al., 2022). Trap sites began as bait sites, where corn was placed with a camera to assess pig visitation. Then, traps were built at bait sites where pig visitation was observed. We used a systematic spacing of approximately 500 m among trap sites for trapping in an effort to saturate the removal zone and maximize bait access to wild pigs that have small space use areas (Kay et al., 2017), while trying to minimize unnecessary overlap where the same wild pigs used multiple sites (Davis et al., 2017; McRae et al., 2020; Snow and VerCauteren, 2019). At each site, we used motion-activated, non-cellular trail cameras (RECONYX PC900, RECONYX Inc., Holmen, WI, USA) and checked cameras at least twice per week for non-active traps or bait sites to assess wild pig visitation before, during, and after trap deployment. Active traps were checked daily, and pigs were dispatched the morning after capture by ground staff. We utilized a variety of trap types including corral and box traps with remotely activated gates (Jager Pro, LLC., Forston, GA, USA; HogEye, Crawford, MS, USA), and passive net corral traps (PigBrig®, PIG BRIG, Field Engine Wildlife Research & Management, Moodus, CT, USA). Overall, we removed 296 wild pigs via trapping.

### 2.4 Toxic baiting removals with carcass recovery

The toxic bait contained 5% sodium nitrite (SN; HOGGONE®; Animal Control Technologies Australia P/L, Victoria, Australia) and was deployed following methods described in (Snow et al., 2021). The process of pre-baiting and ultimate deployment of toxic bait was conducted throughout 23 days, which included ∼14–19 days of pre-baiting sites and acclimating wild pigs to a wild pig-specific bait station before deploying toxic bait at 11 bait sites for 1–2 nights. We deployed 2 nights of non-toxic bait following toxic baiting, and if any surviving wild pigs were observed we deployed another 1-2 nights of toxic bait. In line with ASF introduction response recommendations to remove carcasses of culled wild pigs (United States Department of Agriculture, 2023), we conducted 2 rounds of 400 x 400 m transect grid searches surrounding the bait sites follow toxic baiting to locate carcasses, and ultimately found 41 dead wild pigs. However, we estimated a total removal of 58 wild pigs with the toxic bait using remote cameras and identifying which wild pigs consumed bait and did not return to bait sites following toxic baiting. The carcasses we did not recover likely succumbed to the toxic bait outside of our transect grids.

### 2.5 Aerial Removals with carcass recovery

We flew three consecutive days in the aerial zone from 03/27/2023-03/29/2023 using two helicopters flown by USDA/APHIS/Wildlife Services. We followed the helicopters with 10 ground-personnel (5 teams of 2 individuals each) to locate carcasses (per Feral Swine playbook recommendations (United States Department of Agriculture, 2023)), which were recovered by navigating to geolocations dispatched by the aerial operations crew by radio. Overall, we removed 256 wild pigs via aerial operations, and recovered 126 (49%) of the carcasses.

### 2.6 After-action assessment

We held discussions among crew members to identify operational challenges present in each removal method in the context of TAD responses in different environments. We identified operational characteristics that improve likelihood of achieving disease freedom (Table 1). Then, we discussed operational challenges that could hinder TAD elimination with respect to these characteristics for each removal technique we used, followed by broader operational challenges as part of an integrative management framework (IMF) using our operational knowledge of the strategies we deployed.

**Table 1.** Four operational characteristics of TAD responses, and justification related to how each characteristic relates to objective of disease elimination, we used to identify operational challenges.

| TAD response operation characteristics | Relationship to TAD elimination objective |
| --- | --- |
| Rapid deployment following a TAD detection | Improved likelihoods for elimination when response is early in outbreak (Pepin et al., 2022; Bozzuto et al., 2020; Domenech et al., 2006) |
| Efficient use of time and resources to achieve disease freedom | TAD responses are often costly (Winter-Nelson and Rich, 2008) and require effort over a long duration (Gortázar et al., 2015) |
| Avoidance of counterproductive outcomes | Behavioral responses to culling in other systems have compromised disease elimination achievement, perturbation hypothesis (e.g., increased *Mycobacterium bovis* spillover to cattle from badgers (*Meles meles*) resulting from behavioral responses to culling)(Carter et al., 2007) |
| Implementation at a population/landscape scale | TAD responses often need to be implemented at a landscape scale response in a control zone in order to have adequate coverage of the TAD-infected wild animal population (e.g., ASF Feral Swine response, United States Department of Agriculture, 2023; Desvaux et al., 2021). |

## 3. After-action assessment results: operational challenges

Here, our goal was to identify common challenges (Table 2) for a set of control methods that are likely to affect the operationalization of a landscape-scale TAD response (Table 1).

**Table 2.** Operational challenges identified for control strategies utilized during our study, with corresponding research needs.

|  | Challenge | Solution |
| --- | --- | --- |
| Trapping/Toxicant | Situational awareness (daily) of bait site status are slow and labor-intensive. | Data analysis pipelines to automatically assimilate camera image information with wild pig visitation/equipment status at bait sites with remote cameras, custom data entry software with an automated reporting backend |
|  | Identifying spatial trap and toxicant-deployment strategies | Field and modeling studies to evaluate alternative spatial trapping designs on time to wild pig attraction and control in a defined spatial area |
|  | Identify temporal deployment strategies (e.g., trap/bait site retirement timing/building/setting) | Analytical methodology in the form of a condition-specific rule-of-thumb or more complex modeling-based tool to make trapping decisions |
| Aerial operations | Deployment is highly sensitive to weather conditions | Develop contingency actions for integrated management actions |
|  | Effectiveness of aerial operations on different age classes/behavioral strategies | Field and modeling studies identifying optimal strategies and value of removing individuals across age/behavior |
| Carcass removal | Carcass detection and disposal is labor-intensive and time-consuming | Field studies to measure time, effectiveness, and resource needs for different carcass removal strategies across context and control methods |
|  |  | Development and validation of methods (e.g., UAS, dogs) to efficiently detect carcasses |
|  | Uncertainty in proportion of carcass removals needed | Modeling to evaluate sensitivity of TAD elimination success to proportions of carcass removals with variable decomposition rates. |
| Integrative management framework | How to optimize mixed management strategies during continuous, high-intensity deployment | Field studies to optimize deployment strategy for mixed management to eliminate disease |
|  |  | Stakeholder engagement to reduce operational implementation barriers |
|  |  | Scenario modeling or field study to optimize response time under logistical delays (e.g., inaccessible lands, equipment delays, weather events) |
|  | Inexperience with intense disease control in wild species | Send personnel out to where disease exists, practicing intense removal in different environments, invite personnel from other countries for feedback |

### 3.1 Trapping and toxic bait TAD preparedness challenges

Operational challenges we identified for trapping and toxic bait TAD control included information continuity of trap/bait site situational awareness, and identifying spatial and temporal deployment strategies that maximize near-complete removal of pigs in a TAD response context. With respect to information continuity of situational awareness, we mean maintaining consistent knowledge of pig presence, and the level of pig presence and habituation, to bait/trap sites to inform management actions across crew members. Operational features that led to this challenge were maintenance of a large number of sites, use of manual cameras to characterize pig visitation to sites, and use of multiple rotating crews. While we used a mobile application to enter data, information continuity was interrupted by insufficient cell reception that prevented automatic synching of visitation data across mobile devices in real time.

The second challenge we identified for trapping and toxicant methodologies was spatial optimization of trap and bait sites (i.e., the spatial placement, configuration, and density of traps or bait sites). Our systematic spacing of ∼500m between trap sites was to saturate the removal zone and ensure that all pigs within the active trapping area were likely to encounter a trap site. Yet, we found that our distribution of bait sites may have been too dense, because we frequently observed situations where wild pigs visited multiple bait sites while avoiding nearby sites with active traps.

Lastly, we identified challenges in optimizing temporal deployment strategies for toxicant and trapping methodologies. One specific temporal deployment challenge was deciding when to transition a site to the next stage (e.g., begin trap construction, set a trap, retire a bait site). While we set *a priori* guidelines specifying conditions to transition a site based on pig visitation, adhering to guidelines was challenging because there was uncertainty over whether an area was truly free of wild pigs, especially as fewer wild pigs were observed on camera traps at bait sites.

### 3.2 Aerial operations TAD preparedness challenges

We observed two main operational challenges during aerial removals. The first challenge we found during aerial operations was variable weather conditions that created uncertainty around aerial operation timing. This uncertainty resulted in logistical coordination difficulties that led to an insufficient carcass-searching crew during the first day of aerial operations, alongside a lower carcass recovery proportion, compared to days with sufficient personnel. The second challenge we found during aerial operations was that piglets were difficult to remove by this method.

### 3.3 Carcass recovery TAD preparedness challenges

We found that locating carcasses is time-consuming and inefficient, and there was uncertainty in the proportion of known carcasses that should be removed and how rapidly they should be removed to allow for TAD elimination. The daily proportion of known carcasses we recovered increased with more personnel, but this process was still time-consuming for both aerial and toxicant operations. We also found that 100% recovery of carcasses is unlikely even in this context where we knew approximate locations of carcasses (compared to carcasses resulting from disease mortalities).

### 3.4 Integrative Management Framework TAD preparedness challenges

During our after-action review, we also discussed potential challenges of implementing a combined management effort. While we did not implement an integrative management framework, we referenced our operational experience with the three deployed removal strategies to inform our discussion. Two challenges we identified were optimization of mixed management strategies (i.e., how to efficiently combine removal strategies, or challenges that would affect efficiency of any removal method), and inexperience with landscape-scale disease control in wild species by wildlife management personnel (i.e., few case studies to draw from).

## 4. Discussion: Priorities for addressing deployment challenges

We observed a number of operational challenges in the context of an ASF response in wild pigs, in an arid landscape with low cover. While certain challenges may be specific to our study site, many of the operational challenges we encountered are generalizable to other landscapes and TAD introductions in that they all relate to operational needs for a successful TAD response (Table 1), regardless of location or disease system. Here, we discuss potential solutions (Table 2) for each of the operational challenges we identified, application to TAD response operations across context, and limitations of our inferences.

### 4.1 Trapping and toxicant removals for TAD responses

Challenges we identified during trapping and toxicant removals were information continuity of geographic situational awareness, and optimization of spatial/temporal trap/bait site strategy. The need for information continuity across groups and throughout time/space is a common operational issue across broader emergency response and operational contexts (Dilmaghani and Rao, 2008; Kripalani et al., 2007) which have been rectified by improved communication infrastructure such as wireless mesh networks (Dilmaghani and Rao, 2008). However, communication solutions for TAD responses in wildlife need to incorporate possible remote location and large geographic operational area. Improved automated processing of camera trap or bait/trap site status data, with wireless networks or satellites (e.g., Whytock et al. (2023)), may address information continuity situational awareness needs in this way. Some AI image processing tools currently exist (Tabak et al., 2019), and these tools run batches of images collected from cameras periodically. Development of automated camera trap pipelines in full are needed, alongside testing best practices for transferring data, since rate of data transfer can also lead to communication bottlenecks (Dilmaghani and Rao, 2008). While we encountered this challenge during wild pig trapping, automated camera trap pipelines are likely to be beneficial for other TAD responses as well, since abundance of a host species is an important metric for monitoring population abundance during TAD control efforts (Norouzzadeh et al., 2021; Norouzzadeh et al., 2018) to determine when to modify or stop control operations (Gortazar et al., 2015).

We also noted inefficiencies in spatial and temporal trapping optimization. Specifically, we noted that bait/trap sites may have been redundant. The problem of spatial optimization of traps is not isolated to trapping wild pigs. In fact, alterations in trapping grid density and configuration has been attributed to differences in capture times and probability in insects (El-Sayed et al., 2006) and small mammals (Pearson and Ruggiero, 2003; Warburton and Gormley 2015), suggesting that optimization of spatial trapping design is a more general problem across taxa. While emergency response operators in all regions may not use trapping to reduce wildlife population densities, it is an important method in regions where other methods like aerial operations are infeasible due to high cover (Davis et al., 2018). Further, optimization of trapping/baiting designs at a landscape scale may be an important consideration outside of culling strategies for TAD control, such as vaccination programs (McClure et al., 2022) where consistent bait delivery across a population is needed. Altogether, additional field and modeling studies are needed to identify optimal baiting/trapping strategies for any wildlife species at risk for carrying a TAD, where bait delivery or trapping of live individuals would be part of a TAD control response.

Lastly, we noted difficulties in decision-making on bait/trap site status during trapping/toxicant deployment, which became pronounced towards the end of our trapping effort when there was uncertainty over whether sites were free of pigs. Greater effort (i.e., more time monitoring traps and trying to capture pigs) is a known consequence of decreases in abundance (Fischer et al., 2020). Fine-scale decisions about when to prioritize trapping in other areas or continue monitoring a particular bait/trap site is likely to be relevant in any TAD control context where operators are reducing population densities of a target wildlife species or vector, and this uncertainty is also likely to exist for disease freedom due to inherently higher uncertainty in estimating prevalence when disease prevalence is low (Lachish and Murray, 2018). A solution to aid this decision-making at low densities could be informed by statistical models borrowed from species eradication or wildlife damage control management contexts that interpret sequential ‘zero detection’ days with relevant covariates to relay a probability of presence of the target species (Davis et al., 2017; Parkes and Panetta, 2009; Ramsey et al., 2009) or disease (Davis et al., 2019; Anderson et al. 2013). Further, developing automated workflows that feed camera data into user-friendly applications with these underlying statistical models could reduce communication overhead, increase operational efficiency, and improve probability of achieving TAD elimination as rapidly as possible.

### 4.2 Aerial removals

For challenges in aerial removals, operational challenges we noted were unreliable weather conditions leading to delays, and difficulty removing piglets. While weather conditions will vary seasonally and across study sites, aerial operations are inherently sensitive to weather conditions due to limitations of flight by helicopters. Given this sensitivity, there may be situations where weather conditions require use of alternate strategies, even in areas where aerial removal strategies may be advantageous overall. As with other emergency responses, it is important to plan for the emergency itself as well as the environment in which it takes place (Thévenaz and Resodihardjo 2009). As such, development of contingency plans, dependent on local climate, may be one solution to plan for this sensitivity. Specifically, this might involve plans to utilize other removal techniques following certain thresholds of wait-times for appropriate weather conditions, or identifying personnel deployment alternatives under alternative schedules. For development of contingency plans at the national level, local and experiential knowledge of pilots in regions across the U.S. where wild pig introductions could occur may be an important resource to anticipate seasonal or spatial limitations to aerial operations (Speirs et al., 2021).

We also found that it was difficult to remove piglets during aerial operations. While the timing of lethal control efforts to mitigate wild pig damage can often be flexible enough to occur during low-birthing periods of the year to avoid piglets (Pepin et al., 2017), TAD responses need to be conducted as soon as a TAD introduction is identified (Pepin et al., 2022), which may overlap with a birth pulse. Aerial control operations are very effective for quickly reducing population density of target species, particularly ungulates, across large geographic areas (Bengsen et al., 2022; Choquenot et al., 1999), and are thus important to consider for any TAD control plan involving culling of ungulates. However, field and modeling studies are needed to quantify selective bias of age classes of aerial operations and determine the value of removing particular age classes with respect to controlling disease spread. In the context of TAD responses, selective removal of adults over younger age classes may have deleterious consequences for disease spread if older members of the population have built immunity (Bolzoni et al., 2007). In fact, this phenomenon may be a driving factor behind counterproductive outcomes of culling for wildlife disease elimination purposes (Bolzoni et al., 2007). Since many TAD outbreaks result in some proportion of adults with immunity (e.g., Classical Swine Fever (CSF)(Bolzoni et al., 2007)), selective bias of age classes during aerial removal strategies needs to be quantified with respect to disease characteristics of TADs of interest in order to estimate epidemiological outcomes.

### 4.3 Carcass removals

In our study, we conducted recoveries of culled carcasses, which is notably a different operation than removal of pigs that died from disease, because we knew the general locations of the carcasses. However, even under this context where the carcass locations were known, we still noted challenges in this process including <100% recovery of carcasses, and that it was a time and labor-intensive process. Our simulated control effort was designed with ASFv control in mind, but improving operations of carcass removal is important for any control effort involving a pathogen with carcass-based transmission (e.g., ASFv, Pepin et al., 2022; Screw worm (*ochliomyia hominivorax*), Skoda et al., 2018).

One solution to address the problem of carcass recovery and removal being inefficient is identification of reasonable objectives (i.e., how many need to be removed to be successful in disease elimination?). The importance of recovery and removal of carcasses resulting from disease mortalities has been demonstrated for pathogens such as ASFv that utilize carcass-based transmission in theoretical models (Gervasi and Gubertì, 2022; Pepin et al., 2020; Guinat et al., 2017; Lange, 2015) and successful international ASFv elimination efforts (Danzetta et al., 2020). However, there is a need for to quantify reasonable objectives for removal of culled carcasses because not all culled carcasses will be infectious. Additionally, rapid decomposition in warmer climates may reduce carcass-based transmission rates (Bradhurst et al. 2021, Bowden et al., 2023), suggesting that carcass recovery and removal may not be uniformly effective for ASF control in all situations. Altogether, research to quantify the value of carcass recovery and removal during high culling scenarios and warmer climates, in comparison to other removal techniques, is needed.

Alternative recovery methods could improve efficiency of recovery (both for carcasses resulting from culling and natural deaths) including use of detection dogs (Desvaux et al., 2021) or drone technologies (i.e., unmanned aircraft systems (UAS)). Unmanned aircraft systems have been used for almost 20 years to conduct aerial surveys of wildlife species (Brown et al., 2023; Dunn et al., 2021; Preston et al., 2021; Brisson-Curadeau et al., 2017; Schofield et al., 2017). Increased flight times, miniaturization of sensors, and decreased costs of UAS have improved the ability of this technology to offer more accurate, precise, and timely wild species survey data than traditional methods (Hodgson et al., 2018), including in epidemiological applications (Poljak and Šterbenc, 2020).

### 4.4 Integrative mgmt. framework

We utilized trapping, aerial control, and toxicant removals to simulate a TAD response to ASF. However, applicability of all possible techniques under a TAD response, as with other kinds of emergency responses and disease control efforts, may vary depending on political (Okello et al. 2015), or regulatory (Wilson and Mcreight 2012) constraints. For example, wild pig toxicant control is currently being evaluated for safety to other wildlife species in the U.S. and Australia (Snow et al. 2017) and as such, is still legally prohibited in many countries (Guberti et al. 2019) Further limitations may result from environmental or ecological constraints. For example, aerial control may be less suitable in situations with high foliage cover, or where wild pig densities are very, in which case trapping may be preferable (Wilson and Gentle 2022). However, technological advancements such as thermal-assisted-shooting may improve utility of this method in these less-optimal situations (Wilson and Gentle 2022).

Aside from these nuances, understanding challenges in implementing combinations of these techniques, and challenges that apply broadly to all removal techniques, may be useful for planning a TAD response in an integrative management framework in situations where all removal techniques are tenable. The first challenge we identified for integrative management in TAD responses was optimizing a combined strategy in a TAD elimination context. One consideration for optimizing a TAD response is to measure behavioral changes of target species during TAD control efforts. While culling can be necessary and successful in certain contexts (Prentice et al. 2019), culling actions can also also result in counterproductive disease transmission outcomes (Carter et al. 2007). For example, for badger (*Meles meles*) reservoirs of bovine Tb (*Mycobacterium bovis*), culling actions can result in increased spillover incidents of *M. bovis* to domestic cattle (Carter et al., 2007). For this system, these counterproductive outcomes are likely the result of social perturbations, but the extent of this effect on spillover varies across space and intensity of effort (Carter et al. 2007). In wild pigs, low intensity control operations may have none to minimal effects on pig movement (Bastille-Rousseau et al., 2021; Campbell et al., 2012; Fischer et al., 2016; Campbell et al., 2010) and contact (Yang et al., 2021), but it is unknown whether pig movements would increase under more continuous or intense landscape-scale operations characteristic of TAD response operations (Pepin et al., 2022).

A second solution for optimizing an integrative management framework for TAD responses is to improve efficiency of containment fencing. Although fencing can be effective in controlling movements of wild pigs even under increased pressures of aerial control operations (Lavelle et al., 2011) and has been used with success during some TAD control operations (Chenais et al. 2024), this method has not been uniformly effective in TAD control (Rossi et al. 2015, Han et al. 2022). Operationally, fencing large areas is expensive, labor intensive, and the time required to erect fences may allow disease to spread before completion (Rossi et al. 2015). As such, there is merit to exploration of alternative barriers that are less expensive and easier to install, such as netting. Improving efficiency of fencing options is likely to benefit any TAD control operation where movement wildlife is likely to spread disease and counter TAD elimination efforts.

A third solution to optimize TAD integrative management frameworks is stakeholder engagement, because time for communication and obtaining appropriate permissions after an outbreak has occurred could compromise rapid deployment. For example, during a FMD response in 2007, legal challenges associated with accessing private land resulted in operational delays that reduced the efficacy of the response, suggesting this type of concern could pose challenges to implementing a TAD response elsewhere (Great Britain: Department for Environment, Food and Rural Affairs, 2008). How best to implement stakeholder engagement for TAD responses is beyond the scope of this article, but inefficiencies can arise from differences in values and risk perception in tandem with poor communication and lack of two-way dialogue (Anthony, 2004; Carter et al., 2021). To reduce implementation barriers of control operations during a TAD response, stakeholder engagement ought to be done as soon as possible (ideally *a priori*) to allow time for two-way communication— a critical component of stakeholder engagement (Anthony, 2004; Carter et al., 2021). One method to decision-making that incorporates stakeholder values is a SDM (structured decision model), which not only makes value judgements explicit, but allows for value-based deliberations that incorporate tradeoffs and uncertainties (United States Geological Survey, 2018). Applied to a TAD wild species control operations context, a SDM could incorporate values, tradeoffs, and uncertainties between different control strategies as relevant to stakeholders such as landowners, hunters, large- and small-scale farmers, collaborating local and national governmental entities, and members of other publics.

To address the challenge of personnel inexperience with TAD responses, training exercises developed for TAD control in domestic animals are useful. They provide self-measured preparedness and lead to preparedness-informed actions of participating farmers and producers (Linskens et al., 2018). However, while table-top exercises provide some important strategic planning training, there is no substitute for field-based practice when it comes to identifying potential deployment challenges. Thus, field training for personnel in organizations that may aid wild species population control efforts for a TAD response, including key stakeholders, led by professionals with direct experience controlling wild species for TAD elimination, may be beneficial in improving operational preparedness and may improve coordination among relevant agencies. Such training for TAD preparedness may best be delivered by sending personnel to where TAD elimination is ongoing to practice intense control in different environments.

## 5. Conclusion

Given the unique nature of intensive control to eliminate a TAD in a wild animal population, we identified a number of operational challenges across a discrete set of control methods we utilized. Regardless, there are a number of ways the challenges we identified are applicable to TAD control of other widespread wild animal species and disease systems (e.g. rodents and monkeypox, deer and foot-and-mouth disease, etc.). Given that TAD responses frequently have high costs (Winter-Nelson and Rich, 2008), finding ways to utilize resources in an efficient and cost-effective way can reduce risk of budgetary limitations, and improve TAD response capacity in resource-poor countries, which already face heightened barriers to TAD elimination (Yadav et al., 2020) as well as increased societal consequences of animal disease outbreaks (Goutard et al., 2015). Lastly, understanding reasonable objectives and counterproductive actions of management actions at a landscape scale can reduce time to substantiating disease freedom and avoid unintended ecological consequences.

Altogether, the specifics of how to eliminate TADs in alternative systems may vary depending on the ecology of the wild animal species (i.e. movement patterns, sociality) (Pepin et al., 2022) and the epidemiology of pathogen (i.e., transmission modes, persistence in the environment, host specificity)(Caley and Hone, 2004; Smith et al., 2005). However, we expect that wildlife disease managers will be able to gauge the importance of each of the operational challenges we identified here using knowledge of their disease system, host species, local environment, and chosen removal methods. Given increased rates of pathogen introductions worldwide with increasing global connectivity (Baker et al., 2022), research to identify and address operational challenges of TAD response efforts in different wild animal systems and environments will continue to be a high priority need.

## 6. Acknowledgements

We thank H. Ownbey and G. Studdard for access to private property. We thank Animal Control Technologies Australia for providing the placebo and SN-toxic bait. We thank multiple people from Texas Wildlife Services and Texas Parks and Wildlife for assisting with data collection. We thank anonymous reviewers for their comments on this manuscript.

## References

Anderson, D.P., Ramsey, D.S.L., Nugent, G., Bosson, M., Livingstone, P., Martin, P. a. J., Sergeant, E., Gormley, A.M., Warburton, B., 2013. A novel approach to assess the probability of disease eradication from a wild-animal reservoir host. Epidemiology & Infection 141, 1509–1521. 10.1017/S095026881200310X

Anthony, R., 2004. Risk communication, value judgments, and the public-policy maker relationship in a climate of public sensitivity toward animals: revisiting britain’s foot and mouth crisis. Journal of Agricultural and Environmental Ethics 17, 363–383. 10.1007/s10806-004-5187-2

Bailey, R., 1998. Ecoregions, the ecosystem geography of the oceans and continents. Bailey, R., 1980. Description of the ecoregions of the United States.

Baker, R.E., Mahmud, A.S., Miller, I.F., Rajeev, M., Rasambainarivo, F., Rice, B.L., Takahashi, S., Tatem, A.J., Wagner, C.E., Wang, L.-F., Wesolowski, A., Metcalf, C.J.E., 2022. Infectious disease in an era of global change. Nat Rev Microbiol 20, 193–205. 10.1038/s41579-021-00639-z

Bastille-Rousseau, G., Schlichting, P.E., Keiter, D.A., Smith, J.B., Kilgo, J.C., Wittemyer, G., Vercauteren, K.C., Beasley, J.C., Pepin, K.M., 2021. Multi-level movement response of invasive wild pigs (*Sus scrofa*) to removal. Pest Management Science 77, 85–95. 10.1002/ps.6029

Bengis, R.G., Kock, R.A., Fischer, J., 2002. Infectious animal diseases: the wildlife/livestock interface. Rev Sci Tech 21, 53–65. 10.20506/rst.21.1.1322

Bengsen, A.J., Forsyth, D.M., Pople, A., Brennan, M., Amos, M., Leeson, M., Cox, T.E., Gray, B., Orgill, O., Hampton, J.O., Crittle, T., Haebich, K., 2022. Effectiveness and costs of helicopter-based shooting of deer. Wildl. Res. 50, 617–631. 10.1071/WR21156

Bolzoni, L., Real, L., Leo, G.D., 2007. Transmission Heterogeneity and Control Strategies for Infectious Disease Emergence. PLOS ONE 2, e747. 10.1371/journal.pone.0000747

Bolzoni, L., Tessoni, V., Groppi, M., De Leo, G.A., 2014. React or wait: which optimal culling strategy to control infectious diseases in wildlife. J. Math. Biol. 69, 1001– 1025. 10.1007/s00285-013-0726-y

Bowden, C.F., Grinolds, J., Franckowiak, G., McCallister, L., Halseth, J., Cleland, M., Guerrant, T., Bodenchuk, M., Miknis, R., Marlow, M.C., Brown, V.R., 2023. Evaluation of the effect of hydrated lime on the scavenging of feral swine (S*us scrofa*) carcasses and implications for managing carcass-based transmission of African swine fever virus. Journal of Wildlife Diseases 59, 49–60. 10.7589/JWD-D-22-00061

Bozzuto, C., Schmidt, B.R., Canessa, S., 2020. Active responses to outbreaks of infectious wildlife diseases: objectives, strategies and constraints determine feasibility and success. Proceedings of the Royal Society B: Biological Sciences 287, 20202475. 10.1098/rspb.2020.2475

Bradhurst, R., Garner, G., Roche, S., Iglesias, R., Kung, N., Robinson, B., Willis, S., Cozens, M., Richards, K., Cowled, B., Oberin, M., Tharle, C., Firestone, S., Stevenson, M., 2021. Modelling the spread and control of African swine fever in domestic and feral pigs. URL https://cebra.unimelb.edu.au/data/assets/pdf_file/0003/3949500/CEBRA_project_20121501_final_report_1.4.pdf.

Brisson-Curadeau, É., Bird, D., Burke, C., Fifield, D.A., Pace, P., Sherley, R.B., Elliott, K.H., 2017. Seabird species vary in behavioural response to drone census. Sci Rep 7, 17884. 10.1038/s41598-017-18202-3

Brown, A.M., Allen, S.J., Kelly, N., Hodgson, A.J., 2023. Using Unoccupied Aerial Vehicles to estimate availability and group size error for aerial surveys of coastal dolphins. Remote Sensing in Ecology and Conservation 9, 340–353. 10.1002/rse2.313

Caley, P., Hone, J., 2004. Disease transmission between and within species, and the implications for disease control. Journal of Applied Ecology 41, 94–104. 10.1111/j.1365-2664.2004.00867.x

Campbell, T.A., Long, D.B., Lavelle, M.J., Leland, B.R., Blankenship, T.L., Vercauteren, K.C., 2012. Impact of baiting on feral swine behavior in the presence of culling activities. Prev Vet Med 104, 249–257. 10.1016/j.prevetmed.2012.01.001

Campbell, T.A., Long, D.B., Leland, B.R., 2010. Feral Swine Behavior Relative to Aerial Gunning in Southern Texas. The Journal of Wildlife Management 74, 337–341. 10.2193/2009-131

Carter, L., Mankad, A., Zhang, A., Curnock, M.I., Pollard, C.R.J., 2021. A multidimensional framework to inform stakeholder engagement in the science and management of invasive and pest animal species. Biol Invasions 23, 625–640. 10.1007/s10530-020-02391-6

Carter, S.P., Delahay, R.J., Smith, G.C., Macdonald, D.W., Riordan, P., Etherington, T.R., Pimley, E.R., Walker, N.J., Cheeseman, C.L., 2007. Culling-induced social perturbation in Eurasian badgers *Meles meles* and the management of TB in cattle: an analysis of a critical problem in applied ecology. Proc Biol Sci 274, 2769–2777. 10.1098/rspb.2007.0998

Chenais, E., Depner, K., Guberti, V., Dietze, K., Viltrop, A., Ståhl, K., 2019. Epidemiological considerations on African swine fever in Europe 2014–2018. Porc Health Manag 5, 6. 10.1186/s40813-018-0109-2

Chenais, E., Ahlberg, V., Andersson, K., Banihashem, F., Björk, L., Cedersmyg, M., Ernholm, L., Frössling, J., Gustafsson, W., Hellqvist Björnerot, L., Hultén, C., Kim, H., Leijon, M., Lindström, A., Liu, L., Nilsson, A., Nöremark, M., Olofsson, K.M., Pettersson, E., Rosendal, T., Sjölund, M., Thurfjell, H., Widgren, S., Wikström-Lassa, E., Zohari, S., Ågren, Erik, Ågren, Estelle, Ståhl, K., 2024. First Outbreak of African Swine Fever in Sweden: Local Epidemiology, Surveillance, and Eradication Strategies. Transboundary and Emerging Diseases 2024, 6071781. 10.1155/2024/6071781

Chinchio, E., Crotta, M., Romeo, C., Drewe, J.A., Guitian, J., Ferrari, N., 2020. Invasive alien species and disease risk: An open challenge in public and animal health. PLOS Pathogens 16, e1008922. 10.1371/journal.ppat.1008922

Choquenot, D., Hone, J., Saunders, G., 1999. Using aspects of predator-prey theory to evaluate helicopter shooting for feral pig control. Wildlife Research 26, 251–261. 10.1071/WR98006

Clemmons, E.A., Alfson, K.J., Dutton, J.W., 2021. Transboundary Animal Diseases, an Overview of 17 Diseases with Potential for Global Spread and Serious Consequences. Animals (Basel) 11, 2039. 10.3390/ani11072039

Danzetta, M.L., Marenzoni, M.L., Iannetti, S., Tizzani, P., Calistri, P., Feliziani, F., 2020. African Swine Fever: Lessons to Learn From Past Eradication Experiences. A Systematic Review. Front Vet Sci 7, 296. 10.3389/fvets.2020.00296

Davis, A.J., Kirby, J.D., Chipman, R.B., Nelson, K.M., Xifara, T., Webb, C.T., Wallace, R., Gilbert, A.T., Pepin, K.M., 2019. Not all surveillance data are created equal—A multi-method dynamic occupancy approach to determine rabies elimination from wildlife. Journal of Applied Ecology 56, 2551–2561. 10.1111/1365-2664.13477

Davis, A.J., Leland, B., Bodenchuk, M., VerCauteren, K.C., Pepin, K.M., 2018. Costs and effectiveness of damage management of an overabundant species (*Sus scrofa*) using aerial gunning. wilr 45, 696–705. 10.1071/WR17170

Davis, A.J., Leland, B., Bodenchuk, M., VerCauteren, K.C., Pepin, K.M., 2017. Estimating population density for disease risk assessment: The importance of understanding the area of influence of traps using wild pigs as an example. Preventive Veterinary Medicine 141, 33–37. 10.1016/j.prevetmed.2017.04.004

Desvaux, S., Urbaniak, C., Petit, T., Chaigneau, P., Gerbier, G., Decors, A., Reveillaud, E., Chollet, J.-Y., Petit, G., Faure, E., Rossi, S., 2021. How to Strengthen Wildlife Surveillance to Support Freedom From Disease: Example of ASF Surveillance in France, at the Border With an Infected Area. Frontiers in Veterinary Science 8.

Dilmaghani, R.B., Rao, R.R., 2008. An Ad Hoc Network Infrastructure: Communication and Information Sharing for Emergency Response, in: 2008 IEEE International Conference on Wireless and Mobile Computing, Networking and Communications. Presented at the 2008 IEEE International Conference on Wireless and Mobile Computing, Networking and Communications (WIMOB), IEEE, Avignon, France, pp. 442–447. 10.1109/WiMob.2008.103

Domenech, J., Lubroth, J., Eddi, C., Martin, V., Roger, F., 2006. Regional and International Approaches on Prevention and Control of Animal Transboundary and Emerging Diseases. Annals of the New York Academy of Sciences 1081, 90–107. 10.1196/annals.1373.010

Dunn, M.J., Adlard, S., Taylor, A.P., Wood, A.G., Trathan, P.N., Ratcliffe, N., 2021. Uncrewed aerial vehicle population survey of three sympatrically breeding seabird species at Signy Island, South Orkney Islands. Polar Biol 44, 717–727. 10.1007/s00300-021-02831-6

El-Sayed, A.M., Suckling, D.M., Wearing, C.H., Byers, J.A., 2006. Potential of Mass Trapping for Long-Term Pest Management and Eradication of Invasive Species. Journal of Economic Entomology 99, 1550–1564. 10.1093/jee/99.5.1550

Fischer, J.W., McMurtry, D., Blass, C.R., Walter, W.D., Beringer, J., VerCauteren, K.C., 2016. Effects of simulated removal activities on movements and space use of feral swine. Eur J Wildl Res 62, 285–292. 10.1007/s10344-016-1000-6

Fischer, J.W., Snow, N.P., Wilson, B.E., Beckerman, S.F., Jacques, C.N., VanNatta, E.H., Kay, S.L., VerCauteren, K.C., 2020. Factors and costs associated with removal of a newly established population of invasive wild pigs in Northern U.S. Sci Rep 10, 11528. 10.1038/s41598-020-68264-z

Gervasi, V., Gubertì, V., 2022. Combining hunting and intensive carcass removal to eradicate African swine fever from wild boar populations. Prev Vet Med 203, 105633. 10.1016/j.prevetmed.2022.105633

Gortázar, C., Diez-Delgado, I., Barasona, J.A., Vicente, J., De La Fuente, J., Boadella, M., 2015. The Wild Side of Disease Control at the Wildlife-Livestock-Human Interface: A Review. Front. Vet. Sci. 1. 10.3389/fvets.2014.00027

Goutard, F.L., Binot, A., Duboz, R., Rasamoelina-Andriamanivo, H., Pedrono, M., Holl, D., Peyre, M.I., Cappelle, J., Chevalier, V., Figuié, M., Molia, S., Roger, F.L., 2015. How to reach the poor? Surveillance in low-income countries, lessons from experiences in Cambodia and Madagascar. Preventive Veterinary Medicine, 2nd International Conference on Animal Health Surveillance (ICAHS) 120, 12–26. 10.1016/j.prevetmed.2015.02.014

Great Britain: Department for Environment, Food and Rural Affairs, 2008. Foot and mouth disease 2007: a review and lessons learned.

Guberti, V., Khomenko, S., Masiulis, M., Kerba, S., 2019. African swine fever in wild boar ecology and biosecurity. FAO Animal Production and Health Manual No. 22. Rome, FAO, OIE, and EC.

Guinat, C., Vergne, T., Jurado-Diaz, C., Sánchez-Vizcaíno, J.M., Dixon, L., Pfeiffer, D.U., 2017. Effectiveness and practicality of control strategies for African swine fever: what do we really know? Veterinary Record 180, 97–97. 10.1136/vr.103992

Han, J.-H., Yoo, D.-S., Pak, S.-I., Kim, E.-T., 2022. Understanding the transmission of African swine fever in wild boars of South Korea: A simulation study for parameter estimation. Transboundary and Emerging Diseases 69, e1101–e1112. 10.1111/tbed.14403

Hodgson, J.C., Mott, R., Baylis, S.M., Pham, T.T., Wotherspoon, S., Kilpatrick, A.D., Raja Segaran, R., Reid, I., Terauds, A., Koh, L.P., 2018. Drones count wildlife more accurately and precisely than humans. Methods in Ecology and Evolution 9, 1160– 1167. 10.1111/2041-210X.12974

Jo, Y.-S., Gortázar, C., 2021. African Swine Fever in wild boar: Assessing interventions in South Korea. Transbound Emerg Dis 68, 2878–2889. 10.1111/tbed.14106

Joseph, M.B., Mihaljevic, J.R., Arellano, A.L., Kueneman, J.G., Preston, D.L., Cross, P.C., Johnson, P.T.J., 2013. Taming wildlife disease: bridging the gap between science and management. Journal of Applied Ecology 50, 702–712. 10.1111/1365-2664.12084

Kay, S.L., Fischer, J.W., Monaghan, A.J., Beasley, J.C., Boughton, R., Campbell, T.A., Cooper, S.M., Ditchkoff, S.S., Hartley, S.B., Kilgo, J.C., Wisely, S.M., Wyckoff, A.C., VerCauteren, K.C., Pepin, K.M., 2017. Quantifying drivers of wild pig movement across multiple spatial and temporal scales. Mov Ecol 5, 14. 10.1186/s40462-017-0105-1

Keiter, D.A., Mayer, J.J., Beasley, J.C., 2016. What is in a “common” name? A call for consistent terminology for nonnative *Sus scrofa*. Wildlife Society Bulletin 40, 384–387. 10.1002/wsb.649

Kripalani, S., LeFevre, F., Phillips, C.O., Williams, M.V., Basaviah, P., Baker, D.W., 2007. Deficits in Communication and Information Transfer Between Hospital-Based and Primary Care Physicians: Implications for Patient Safety and Continuity of Care. JAMA 297, 831–841. 10.1001/jama.297.8.831

Lachish, S., Murray, K.A., 2018. The Certainty of Uncertainty: Potential Sources of Bias and Imprecision in Disease Ecology Studies. Front. Vet. Sci. 5. 10.3389/fvets.2018.00090

Lange, M., 2015. Alternative control strategies against ASF in wild boar populations. EFSA Supporting Publications 12, 843E. 10.2903/sp.efsa.2015.EN-843

Lavelle, M.J., Vercauteren, K.C., Hefley, T.J., Phillips, G.E., Hygnstrom, S.E., Long, D.B., Fischer, J.W., Swafford, S.R., Campbell, T.A., 2011. Evaluation of fences for containing feral swine under simulated depopulation conditions. The Journal of Wildlife Management 75, 1200–1208. 10.1002/jwmg.134

Lewis, A.A., Williams, B.L., Smith, M.D., Ditchkoff, S.S., 2022. Shifting to sounders: Whole sounder removal eliminates wild pigs. Wildlife Society Bulletin 46, e1260. 10.1002/wsb.1260

Lewis, J.S., Corn, J.L., Mayer, J.J., Jordan, T.R., Farnsworth, M.L., Burdett, C.L., VerCauteren, K.C., Sweeney, S.J., Miller, R.S., 2019. Historical, current, and potential population size estimates of invasive wild pigs (*Sus scrofa*) in the United States. Biol Invasions 21, 2373–2384. 10.1007/s10530-019-01983-1

Licoppe, A., De Waele, V., Malengreaux, C., Paternostre, J., Van Goethem, A., Desmecht, D., Herman, M., Linden, A., 2023. Management of a Focal Introduction of ASF Virus in Wild Boar: The Belgian Experience. Pathogens 12, 152. 10.3390/pathogens12020152

Linskens, E.J., Neu, A.E., Walz, E.J., Charles, K.M.S., Culhane, M.R., Ssematimba, A., Goldsmith, T.J., Halvorson, D.A., Cardona, C.J., 2018. Preparing for a Foreign Animal Disease Outbreak Using a Novel Tabletop Exercise. Prehospital and Disaster Medicine 33, 640–646. 10.1017/S1049023X18000717

McClure, K.M., Bastille-Rousseau, G., Davis, A.J., Stengel, C.A., Nelson, K.M., Chipman, R.B., Wittemyer, G., Abdo, Z., Gilbert, A.T., Pepin, K.M., 2022. Accounting for animal movement improves vaccination strategies against wildlife disease in heterogeneous landscapes. Ecological Applications 32, e2568. 10.1002/eap.2568

McRae, J.E., Schlichting, P.E., Snow, N.P., Davis, A.J., VerCauteren, K.C., Kilgo, J.C., Keiter, D.A., Beasley, J.C., Pepin, K.M., 2020. Factors Affecting Bait Site Visitation: Area of Influence of Baits. Wildlife Society Bulletin 44, 362–371. 10.1002/wsb.1074

Meng, X.J., Lindsay, D.S., Sriranganathan, N., 2009. Wild boars as sources for infectious diseases in livestock and humans. Philos Trans R Soc Lond B Biol Sci 364, 2697– 2707. 10.1098/rstb.2009.0086

Miller, R.S., Sweeney, S.J., Slootmaker, C., Grear, D.A., Di Salvo, P.A., Kiser, D., Shwiff, S.A., 2017. Cross-species transmission potential between wild pigs, livestock, poultry, wildlife, and humans: implications for disease risk management in North America. Sci Rep 7, 7821. 10.1038/s41598-017-07336-z

NOAA, 2023. Climate Data Online [WWW Document]. URL https://www.ncdc.noaa.gov/cdo-web/ (accessed 1.24.24).

Norouzzadeh, M.S., Morris, D., Beery, S., Joshi, N., Jojic, N., Clune, J., 2021. A deep active learning system for species identification and counting in camera trap images. Methods in Ecology and Evolution 12, 150–161. 10.1111/2041-210X.13504

Norouzzadeh, M.S., Nguyen, A., Kosmala, M., Swanson, A., Palmer, M.S., Packer, C., Clune, J., 2018. Automatically identifying, counting, and describing wild animals in camera-trap images with deep learning. Proceedings of the National Academy of Sciences 115, E5716–E5725. 10.1073/pnas.1719367115

Nugent, G., Buddle, B., Knowles, G., 2015. Epidemiology and control of *Mycobacterium bovis* infection in brushtail possums (*Trichosurus vulpecula*), the primary wildlife host of bovine tuberculosis in New Zealand. N Z Vet J 63, 28–41. 10.1080/00480169.2014.963791

Okello, A., Welburn, S., Smith, J., 2015. Crossing institutional boundaries: mapping the policy process for improved control of endemic and neglected zoonoses in sub-Saharan Africa. Health Policy and Planning 30, 804–812. 10.1093/heapol/czu059

Paarlberg, P.L., Hillberg, A., Lee, J.G., Mathews, K.H. (Eds.), 2008. Economic Impacts of Foreign Animal Disease, Economic Research Report. 10.22004/ag.econ.56453

Parkes, J., Panetta, F., 2009. Eradication of invasive species: progress and emerging issues in the 21st century. Invasive species management. 10.1093/oso/9780199216321.003.0004

Pearson, D.E., Ruggiero, L.F., 2003. Transect versus Grid Trapping Arrangements for Sampling Small-Mammal Communities. Wildlife Society Bulletin (1973-2006) 31, 454–459.

Pepin, K.M., Brown, V.R., Yang, A., Beasley, J.C., Boughton, R., VerCauteren, K.C., Miller, R.S., Bevins, S.N., 2022. Optimising response to an introduction of African swine fever in wild pigs. Transboundary and Emerging Diseases 69, e3111–e3127. 10.1111/tbed.14668

Pepin, K.M., Davis, A.J., VerCauteren, K.C., 2017. Efficiency of different spatial and temporal strategies for reducing vertebrate pest populations. Ecological Modelling 365, 106–118. 10.1016/j.ecolmodel.2017.10.005

Pepin, K.M., Golnar, A.J., Abdo, Z., Podgórski, T., 2020. Ecological drivers of African swine fever virus persistence in wild boar populations: Insight for control. Ecology and Evolution 10, 2846–2859. 10.1002/ece3.6100

Pepin, K.M., VerCauteren, K.C., 2016. Disease-emergence dynamics and control in a socially-structured wildlife species. Sci Rep 6, 25150. 10.1038/srep25150

Poljak, M., Šterbenc, A., 2020. Use of drones in clinical microbiology and infectious diseases: current status, challenges and barriers. Clin Microbiol Infect 26, 425–430. 10.1016/j.cmi.2019.09.014

Potapov, A., Merrill, E., Lewis, M.A., 2012. Wildlife disease elimination and density dependence. Proceedings of the Royal Society B: Biological Sciences 279, 3139– 3145. 10.1098/rspb.2012.0520

Prentice, J.C., Fox, N.J., Hutchings, M.R., White, P.C.L., Davidson, R.S., Marion, G., 2019. When to kill a cull: factors affecting the success of culling wildlife for disease control. Journal of The Royal Society Interface 16, 20180901. 10.1098/rsif.2018.0901

Preston, T.M., Wildhaber, M.L., Green, N.S., Albers, J.L., Debenedetto, G.P., 2021. Enumerating White-Tailed Deer Using Unmanned Aerial Vehicles. Wildlife Society Bulletin 45, 97–108. 10.1002/wsb.1149

Ramsey, D.S.L., Parkes, J., Morrison, S.A., 2009. Quantifying Eradication Success: the Removal of Feral Pigs from Santa Cruz Island, California. Conservation Biology 23, 449–459. 10.1111/j.1523-1739.2008.01119.x

Risch, D.R., Ringma, J., Price, M.R., 2021. The global impact of wild pigs (*Sus scrofa*) on terrestrial biodiversity. Sci Rep 11, 13256. 10.1038/s41598-021-92691-1

Rossi, S., Staubach, C., Blome, S., Guberti, V., Thulke, H.-H., Vos, A., Koenen, F., Le Potier, M.-F., 2015. Controlling of CSFV in European wild boar using oral vaccination: a review. Front. Microbiol. 6. 10.3389/fmicb.2015.01141

Ruiz-Fons, F., Segalés, J., Gortázar, C., 2008. A review of viral diseases of the European wild boar: Effects of population dynamics and reservoir rôle. The Veterinary Journal 176, 158–169. 10.1016/j.tvjl.2007.02.017

Schofield, G., Katselidis, K.A., Lilley, M.K.S., Reina, R.D., Hays, G.C., 2017. Detecting elusive aspects of wildlife ecology using drones: New insights on the mating dynamics and operational sex ratios of sea turtles. Functional Ecology 31, 2310– 2319. 10.1111/1365-2435.12930

Skoda, S.R., Phillips, P.L., Welch, J.B., 2018. Screwworm (*Diptera: Calliphoridae*) in the United States: Response to and Elimination of the 2016–2017 Outbreak in Florida. Journal of Medical Entomology 55, 777–786. 10.1093/jme/tjy049

Smith, K.F., Dobson, A.P., McKenzie, F.E., Real, L.A., Smith, D.L., Wilson, M.L., 2005. Ecological theory to enhance infectious disease control and public health policy. Frontiers in Ecology and the Environment 3, 29–37. 10.1890/1540-9295(2005)003[0029:ETTEID]2.0.CO;2

Snow, N.P., Foster, J.A., Kinsey, J.C., Humphrys, S.T., Staples, L.D., Hewitt, D.G., Vercauteren, K.C., 2017. Development of toxic bait to control invasive wild pigs and reduce damage. Wildlife Society Bulletin 41, 256–263. 10.1002/wsb.775

Snow, N.P., VerCauteren, K.C., 2019. Movement responses inform effectiveness and consequences of baiting wild pigs for population control. Crop Protection 124, 104835. 10.1016/j.cropro.2019.05.029

Snow, N.P., Wishart, J.D., Foster, J.A., Staples, L.D., VerCauteren, K.C., 2021. Efficacy and risks from a modified sodium nitrite toxic bait for wild pigs. Pest Manag Sci 77, 1616–1625. 10.1002/ps.6180

Snow, N.P., Smith, B., Lavelle, M. J., Glow, M. P., Chalkowski, K., Leland, B. R., Sherburne, S., Fischer, J. W., Kohen, K. J., Cook, S. M., Smith, H., VerCauteren, K. C., Miller, R. S., Pepin, K. M. Comparing efficiencies of population control methods for responding to foreign animal disease threats in wild pigs. [This work will be available as a preprint prior to publication of the present manuscript, and will be cited directly]

Sokolow, S.H., Nova, N., Pepin, K.M., Peel, A.J., Pulliam, J.R.C., Manlove, K., Cross, P.C., Becker, D.J., Plowright, R.K., McCallum, H., De Leo, G.A., 2019. Ecological interventions to prevent and manage zoonotic pathogen spillover. Philosophical Transactions of the Royal Society B: Biological Sciences 374, 20180342. 10.1098/rstb.2018.0342

Speirs, A., Ramée, C., Payan, A.P., Mavris, D., Feigh, K.M., 2021. Impact of Adverse Weather on Commercial Helicopter Pilot Decision-Making and Standard Operating Procedures, in: AIAA AVIATION 2021 FORUM. American Institute of Aeronautics and Astronautics. 10.2514/6.2021-2771

Steele, L., Orefuwa, E., Dickmann, P., 2016. Drivers of earlier infectious disease outbreak detection: a systematic literature review. International Journal of Infectious Diseases 53, 15–20. 10.1016/j.ijid.2016.10.005

Tabak, M.A., Norouzzadeh, M.S., Wolfson, D.W., Sweeney, S.J., Vercauteren, K.C., Snow, N.P., Halseth, J.M., Di Salvo, P.A., Lewis, J.S., White, M.D., Teton, B., Beasley, J.C., Schlichting, P.E., Boughton, R.K., Wight, B., Newkirk, E.S., Ivan, J.S., Odell, E.A., Brook, R.K., Lukacs, P.M., Moeller, A.K., Mandeville, E.G., Clune, J., Miller, R.S., 2019. Machine learning to classify animal species in camera trap images: Applications in ecology. Methods in Ecology and Evolution 10, 585–590. 10.1111/2041-210X.13120

Thévenaz, C., Resodihardjo, S.L., 2010. All the best laid plans…conditions impeding proper emergency response. International Journal of Production Economics 126, 7–21. 10.1016/j.ijpe.2009.09.009

United States Department of Agriculture, 2023. African Swine Fever Response, Outbreak in Feral Swine: Incident Playbook.

United States Department of Agriculture, 2020. African swine fever response plan: The Red Book.

United States Geological Survey, 2018. Structured Decision Making.

VerCauteren, K.C., Pepin, K.M., Cook, S.M., McKee, S., Pagels, A., Kohen, K.J., Messer, I.A., Glow, M.P., Snow, N.P., 2024. What is known, unknown, and needed to be known about damage caused by wild pigs. Biol Invasions 26, 1313–1325. 10.1007/s10530-024-03263-z

Warburton, B., Gormley, A.M., 2015. Optimising the Application of Multiple-Capture Traps for Invasive Species Management Using Spatial Simulation. PLOS ONE 10, e0120373. 10.1371/journal.pone.0120373

Whytock, R.C., Suijten, T., van Deursen, T., Świeżewski, J., Mermiaghe, H., Madamba, N., Mouckoumou, N., Zwerts, J.A., Pambo, A.F.K., Bahaa-el-din, L., Brittain, S., Cardoso, A.W., Henschel, P., Lehmann, D., Momboua, B.R., Makaga, L., Orbell, C., White, L.J.T., Iponga, D.M., Abernethy, K.A., 2023. Real-time alerts from AI-enabled camera traps using the Iridium satellite network: A case-study in Gabon, Central Africa. Methods in Ecology and Evolution 14, 867–874. 10.1111/2041-210X.14036

Wilson, C., Gentile, M.N., 2022. Feral pig population control techniques: A review and discussion of efficacy and efficiency for application in Queensland. Technical Report. State of Queensland, Brisbane.

Wilson, L., Mccreight, R., 2012. Public Emergency Laws & Regulations: Understanding Constraints & Opportunities. Journal of Homeland Security and Emergency Management 9. 10.1515/1547-7355.2034

Winter-Nelson, A., Rich, K.M., 2008. Mad Cows and Sick Birds: Financing International Responses to Animal Disease in Developing Countries. Development Policy Review 26, 211–226. 10.1111/j.1467-7679.2008.00406.x

Yadav, M.P., Singh, R.K., Malik, Y.S., 2020. Emerging and Transboundary Animal Viral Diseases: Perspectives and Preparedness. Emerging and Transboundary Animal Viruses 1–25. 10.1007/978-981-15-0402-0_1

Yang, A., Schlichting, P., Wight, B., Anderson, W.M., Chinn, S.M., Wilber, M.Q., Miller, R.S., Beasley, J.C., Boughton, R.K., VerCauteren, K.C., Wittemyer, G., Pepin, K.M., 2021. Effects of social structure and management on risk of disease establishment in wild pigs. Journal of Animal Ecology 90, 820–833. 10.1111/1365-2656.13412

